# SYSTEM XC- AS A MOLECULAR MECHANISM FOR EVOLUTIONARY NEW FORMS OF ADVANCED COGNITION

**DOI:** 10.1101/2025.03.17.643792

**Authors:** Gregory Simandl, Robert C. Twining, Nicholas J. Raddatz, Gracie Berentson, Sidney Peck, Robert Wheeler, Iaroslav Savtchouk, SuJean Choi, David A. Baker

## Abstract

Human cognitive abilities are deeply rooted in evolutionary building blocks that maximize computation while maintaining efficiency. These abilities are not without evolutionary signatures; conserved processes like vision have undergone continual phylogenetic adjustments to better serve ecological niches. Conversely, more sophisticated forms of cognition may have required evolutionary innovations to transform existing neuronal processing to expand computational abilities. One such innovation is system xc- (Sxc), a cystine-glutamate antiporter predominantly localized to astrocytes that emerged in deuterostomes (e.g., vertebrates) after their divergence from protostomes over 550 million years ago. Previous evidence suggests that genetically modified rats that lack functional Sxc (MSxc) exhibit enhanced cocaine-seeking behavior. In this study, we deconstructed drug-seeking into its component behaviors, categorizing them as reliant on evolutionary conserved or newly evolved cognitive processes. Our results reveal that Sxc function is dispensable for conserved processes like visual, emotional, and hedonic processing, but critical for advanced, evolutionary new cognitive functions, particularly impulse control and decision making. Notably, we demonstrate a temporally specific reliance on Sxc during the learning phase of optimal decision-making, but not in maintaining established strategies. This is an important addition to our current understanding of astrocytes in non-homeostatic functions, indicating their critical role in computationally demanding phases of learning and memory. Unraveling evolutionary innovations like Sxc not only deepens our understanding of cognitive evolution but also paves the way for revolutionary, precision-targeted therapies in neuropsychiatric disorders, potentially transforming treatment paradigms and patient outcomes.

## Introduction

Developing effective therapeutics for neuropsychiatric disorders is hindered by the brain’s remarkable efficiency, which achieves complex cognitive function through a surprisingly limited number of neurotransmitter systems. This efficiency is rooted in molecular degeneracy (Edelman and Gally (2001); Noppeney et al. (2004); Marder (2012); Cropper et al. (2016)), where structurally distinct mechanisms produce similar functional outcomes. This contrasts with molecular redundancy, which merely duplicates identical components for backup purposes. Crucial for evolutionary success, degeneracy enables individual variability and resilience against environmental uncertainties (Albantakis et al. (2024)), permitting multiple degenerate circuits involved in the same function to diverge and adopt novel functions without compromising their original roles (Edelman and Gally (2001); Friston and Price (2003); Barron et al. (2023)).

Neurotransmitter systems exemplify this principle, as structurally diverse molecules can trigger similar cellular responses while serving unique functions in different contexts. For example, the neurotransmitter glutamate has evolved to regulate various forms of brain functions, ranging from visual processing to memory formation, through multiple receptor systems. While degeneracy is an essential process that enables dramatic gains in efficient signaling complexity, the individual variability in signaling network development creates diverse sources of dysfunction that ultimately converge to produce similar clinical outcomes (Stöber et al. (2023); Paunova et al. (2023)). As a result, consensus molecular mechanisms targeted for therapeutic interventions will often result in a wide array of altered brain functions, including off-target adverse effects. A promising approach to addressing these therapeutic challenges is to investigate how phylogenetically advanced traits, such as human cognition, evolved from conserved degenerate systems, leveraging molecular degeneracy’s dual role in maintaining core functions while enabling evolutionary innovation.

Cocaine use disorders (CUDs) are often characterized by persistent cognitive deficits, particularly within domains of impulse control and decision-making, which significantly contribute to increased relapse vulnerability (Bolla et al. (2003); Perry and Carroll (2008); Belin et al. (2008)). These impairments are linked to the dysregulation of glutamate signaling, which exerts critical control over cognitive circuits involved in inhibitory processes, such as impulse control (Piszczek et al. (2022); Carollo et al. (2024)). While dysfunction of glutamate signaling has been strongly linked to CUD (Baker et al. (2003); McFarland et al. (2003); Kalivas and O’Brien (2008); Spencer et al. (2016)), its importance to virtually every circuit in the brain makes it an ideal candidate for elucidating how conserved molecular mechanisms have been repurposed to support complex cognitive functions that are disrupted in addiction.

System xc- (Sxc) represents an evolutionary recent addition to the glutamate signaling network found across all vertebrate species (Lewerenz et al. (2013)). While it is a glutamate-release mechanism expressed by astrocytes throughout the central nervous system, disrupting Sxc-mediated glutamate release in rats is not fatal and does not induce widespread impairments to brain function (Hess et al. (2023)). Notably, repeated cocaine self-administration in rats impairs Sxc function, which contributes to drug seeking behavior (Baker et al. (2002); Baker et al. (2003)). In support, rats genetically modified to eliminate Sxc function (MSxc) exhibit enhanced drug seeking behavior in a cocaine reinstatement paradigm (Hess et al. (2023)).

The purpose of these studies was to gain insight into the discrete cognitive processes requiring intact Sxc function. Specifically, we dissected the complex behavior of drug-seeking into its constitutive parts to uncover how each element contributes to deficits observed in MSxc rats. By categorizing these behaviors as evolutionary conserved (sensory, emotional, and hedonic processing) and recently evolved cognitive functions (impulse control and decision making), these studies may illustrate how the evolutionary emergence of Sxc enhanced cognitive abilities independent of highly degenerate, conserved neural systems.

## Materials and Methods

### Animal care and usage

Male Sprague Dawley rats (post-natal day 70-150) were used in these studies. Rats used for genetic studies were obtained from an internal colony of genetically modified rats generated using a Het/Het breeding approach, which allowed for the use of WT and mutant Sxc (MSxc) littermates in these studies. Outbreeding involved the use of Sprague Dawley rats purchased from Envigo, which occurred every three generations. Rats used for pharmacological studies were obtained from Envigo. After postnatal day 60, rats were individually housed in a temperature and humidity-controlled room, maintained on a 12:12 light/dark cycle with lights off at 0700. Behavioral experiments occurred during the dark cycle, unless otherwise stated. All procedures and protocols were approved by the Institutional Animal Care and Use Committee at Marquette University and adhered to the guidelines set forth by the National Institutes of Health.

### Mutant Rat

For creation of MSxc genetically modified rats, zinc-finger nucleases (ZFNs) were designed to target the second exon sequence and produce small deletions of a limited number of base pairs in the *Slc7a11* gene resulting in whole-animal disruption of *Slc7a11* (MSxc rats). See Hess et al. (2023) for detailed methods.

### Two-Meal Paradigm

#### Diets

Teklad standard chow (SC; 18% protein, 44% carbohydrate, 6% fat; 3.1 kcal/g) or palatable western diet (WD; D12079B; Research Diets; New Brunswick, NJ; 17% protein, 43% carbohydrate, 41% fat, 4.7 kcal/g) (Hurley et al. (2016)).

#### Procedure

Rats (N=14/genotype) were habituated to consume their daily SC intake in a single 2-hour period after the onset of dark phase (Meal 1, M1). After establishing stable feeding and weight gain patterns (30 – 50 kCal/2h; body weight <1% change/day), rats were offered a short 15-minute meal (Meal 2, M2) of either SC or WD (N = 6-8/treatment/genotype) approximately 30 minutes following the removal of M1 for 10 days (Hurley et al. (2016)). Feeding was measured via the BIoDAQ system or by weighing food bins before and after experimental sessions (including spilled food). The BIoDAQ feeding system automates data acquisition and records food intake using algorithmic load cell technology (Research Diets, New Brunswick, JM). Food intake and body weight measurements were recorded, and final session values were used for diet intake comparisons.

### Elevated Plus Maze (EPM)

Rats (N=20-22/genotype) were placed in a plus maze of one cm thick black Plexiglas and elevated at a height of 55 cm from the floor. Two open arms (50.8 cm x 10.2 cm) were connected to two enclosed arms (50.8 cm x 10.2 cm x 30 cm) by an open square (12.7 cm x 12.7 cm). Each rat was placed in the elevated plus maze for five minutes, facing the open arm pointing away from the tester to start. The session was recorded and an observer blind to treatment recorded the number of entries, which was defined when the rat placed all four feet into an open arm. Time of entry in the open arm was recorded from the time the rat placed four feet in the open arm until two of the rat’s feet entered the open square. Behavior was recorded and analyzed by two reviewers blinded to rat genotype.

### Touchscreen Apparatus

Training and testing took place within sound and light-attenuated touchscreen chambers (60cm x 35.2cm x 67cm; Lafayette Instruments). Light stimuli were presented at different positions on the touch screen and sectioned by a task-specific screen mask (Visual Discrimination: 2x1; 5CSRT & rGT: 5x1). Rats learned to associate a light stimulus (illuminated white square) with a food reward (45mg dustless chocolate precision pellets, BioServ). System control was provided by Whisker Standard and ABET II software (Lafayette Instrument).

### Visual Discrimination Task

To examine visual sensory processing through single visual discrimination, rats (N=7-8/genotype) were trained to distinguish positive and negative visual stimuli (CS+ and CS-) assigned to a plane or spider and counterbalanced across subjects. The visual stimulus pairing was selected based on minimal bias for either stimulus (Bussey et al. (2008)). Pretraining and testing consisted of 60 trials across 30 minutes.

#### Pretraining

In the first stage, rats were trained to collect reward pellets delivered under a 30-second variable interval (VI30) paired with the illumination of the food tray light and tone presentation (3 kHz). During this stage, random visual stimuli (varying in shape and brightness) were used to prevent biases and were presented in either of the response windows for 30 seconds. A single pellet was administered for each trial unless the rat touched the stimulus, resulting in a 3-pellet reward. In the next stage, rats were required to respond to the stimulus presented on the touchscreen to obtain a reward, which remained on the screen until selected, establishing a learned behavior of intentional stimulus interaction. In the next stage, rats learned to initiate a trial by engaging with the food tray. In the last stage, rats that selected the incorrect stimulus (blank screen) were punished with a 5-second timeout that resulted in house-light illumination and no reward, followed by a correction period that repeated the same trial conditions until a correct response was made, which was then rewarded.

#### Testing (Single Visual Discrimination)

First, a free reward was delivered with the tray light illuminated. The rat was required to nose-poke and exit the food tray to initiate the first trial. Trials began with two novel stimuli (airplane or spider) presented on the screen, assigned as CS+ and CS-, and positioned pseudo-randomly. Rats learned to associate the CS+ image with reward while avoiding the CS-image. Performance was measured by quantifying the trials to reach criteria (>80% accuracy for two consecutive sessions) and average accuracy percentage on criterion-achieving sessions.

### 5-Choice Serial Reaction Time Task (5CSRT)

To measure impulse control, we utilized the 5-choice serial reaction time task. Initially, rats (N=6-7/genotype) were trained to nose-poke to initiate a trial, associate a light stimulus with a reward (Initial Touch), then nose-poke for the reward (Must Touch). We then reduced the maximum stimulus duration and response window times across successive training schedules (Fig 4B), where a 1 second stimulus duration and 5 second response window was used for testing. If the rat did not respond within the response window (omission), the trial was terminated, and the food tray was illuminated to indicate next trial availability. Rats that engaged with the touch screen before stimulus presentation (premature response) were punished with a 5-second timeout accompanied by house light illumination. Rats were tested across 12 consecutive sessions and final metrics for choice accuracy, omission rate, and premature response rate were recorded on the final testing session (session 12).

### Rat Gambling Task (rGT)

To measure decision making, we employed the rat gambling task. First, rats (N=10/genotype – light phase; N=20/treatment Chronic Baseline – dark phase; N=5-11/treatment Post-Baseline – dark phase) were trained to nose-poke to initiate a trial and associate a light stimulus with a reward (Initial Touch), then nose-poke for the reward (Must Touch). We then reduced the maximum time to nose-poke the light stimulus across successive training schedules (schedule 1: 60 second, schedule 2: 30 second, schedule 3: 15 second, schedule 4: 10 second), where 10 seconds was used for the remainder of the training and testing (Basic Training). If the rat did not respond within the time window (omission), all stimuli were terminated, and the food tray was illuminated to indicate next trial availability. Rats that engaged with the touch screen before stimulus presentation (premature response) were punished with a 5-second timeout accompanied by house light illumination. Next, we trained rats on individual reward/punishment schedules that were spatially assigned to light stimuli P1-4 and presented pseudo-randomly (Fig 5B; Forced Choice). Winning trials resulted in delivery of the chosen stimulus scheduled reward density, while losing trials were accompanied by constant house light illumination and the chosen stimulus light pulsing (0.25Hz) for the scheduled duration. During the testing phase (Free Choice), rats were tested once daily in a 30-minute session where all four light stimuli were presented simultaneously. Rats were tested across 15 days, unless stated otherwise.

#### rGT Pharmacology

To enhance temporal precision of disrupting system xc- (Sxc) function, sulfasalazine (SSZ) was administered either during or after rat gambling free choice baseline behavior was established. SSZ (0-16 mg/kg; IP; Sigma), an inhibitor of Sxc (Bernabucci et al. (2012); Gout et al. (2001); Sontheimer and Bridges (2012); Lutgen et al. (2014)), was dissolved in isotonic saline and brought to a pH between 6.0 and 8.0 using NaOH. Testing began 2 hours after SSZ treatment (Lutgen et al. (2014)). Acute testing involved a single SSZ dose, while chronic testing occurred daily for 12 days during baseline testing or daily for 5 days after baseline testing.

### Experimental Design and Statistical Analysis

Investigators were blinded to treatment (genotype and pharmacology) for all behavioral procedures. All data are presented as means ± SEM. Statistical analyses were conducted using GraphPad Prism (version 10) and Rstudio (GLMM only).

Statistical significance was determined using Fisher’s LSD, Student’s *t* tests, two-way repeated-measures ANOVA, and generalized linear mixed model (GLMM). Fisher’s LSD and Student’s *t* tests were used to make planned pairwise comparisons when there were only two means to compare or when ANOVAs revealed an appropriate significant main effect or interaction. Two-way repeated measures ANOVA varied genotype (MSxc vs WT) and meal (Two-Meal; WD vs SC) as between-subjects factors; and meal (Two-Meal), experimental stage (5CSRT), session (rGT), and timepoint (rGT) as within-subjects factors. The Holms-Sidak procedure was used to contain familywise α level at 0.05 for multiple *post hoc* comparisons. Effects sizes were calculated using partial eta squared (η_p_^2^; ANOVA main effects and interactions) and Cohen’s *d* for important pairwise comparisons (Student’s *t* tests and Fisher LSD comparisons). Unique analytical details for each experiment and follow-up statistical analyses are reported below in Results.

#### Generalized Linear Mixed Model

A generalized linear mixed model (GLMM) was used to analyze the proportion of optimal (P2) choices (P2 choice/Total Choices per session) as an index of performance:

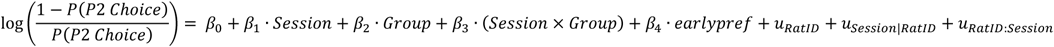

Given the outcome is a proportion, the model was specified with a binomial distribution and the logit link function, modeling the log odds of making a P2 choice. Fixed effects included Group (WT, MSxc), Session (2–15), their interaction (Session × Group), and initial preference (earlypref), categorized from performance on the first session into low (<20%), moderate (20–60%), and high (>60%). This categorization served as a blocking factor to statistically control for potential genotypic biases in rats’ baseline lever preferences. Session 1 data were subsequently excluded from the model to avoid predictor-outcome dependency thus preserving the independence between baseline categorization and subsequent predictive modeling. Random effects (u) for RatID, Session nested within RatID (random slopes), and their interaction RatID × Session accounted for individual differences and repeated measurements, and subject specific learning trajectories. Observations were weighted by the total number of choices per subject per session. Fitted values were transformed to probabilities ranging from 0 to 1 for visualization (Fig 5E & Fig 6B) using the logistic function: 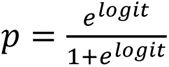

For pharmacological predictions (Fig 6), a similar GLMM framework was applied to analyze the proportion of optimal (P2) choices.

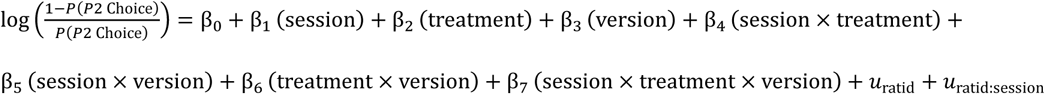

Fixed effects included Session (1–15), Treatment (Vehicle, SSZ), Version (A, B; counterbalancing stimulus positions across subjects), and their interactions (Session × Treatment, Session × Version, Treatment × Version, Session × Treatment × Version). Unlike the genotypic experiment, initial preference was not used as a blocking factor because rats were assigned to treatment groups using a randomized block design based on pre-treatment training performance and treatment administration began on the first test session, thereby preventing systematic pre-existing biases in baseline P2 preference. Random effects (u) were included for RatID and Session nested within RatID (random intercept), accounted for individual differences in baseline choice behavior and session-to-session deviations from individual performance trajectories. A random slope for Session was initially tested but subsequently excluded due to poor model fit with minimal explanatory improvement. Using random intercepts alone effectively accounted for individual variability nested in repeated measures within and across session.

## Results

### Deconstructing Drug Seeking Behavior into Discrete Evolutionary Domains

The complexity of psychiatric disorders like cocaine use disorder (CUD) often hinders effective treatment development. Current approaches frequently result in generalized diagnoses and limited effective therapeutic targets. To address these challenges, we need to 1) better understand complex neuropsychiatric disorders and 2) identify novel molecular targets for specific symptoms. We adopt an evolutionary approach to analyze drug-seeking behavior, a preclinical model for relapse potential in humans (Fig 1). By distinguishing between evolutionarily conserved and newly evolved cognitive circuits, we can pinpoint the specific cognitive domains most vulnerable to relapse. Notably, rats genetically modified to eliminate system xc- function (MSxc) show increased drug-seeking behavior without affecting basic, evolutionarily conserved cognitive processes (Hess et al. (2023)). This positions the MSxc rat as an ideal model for evolutionary cognitive screening and highlights Sxc’s therapeutic potential in addressing phylogenetically recent cognitive deficits associated with various neuropsychiatric disorders (Kaczanowska et al. (2022)).

**Figure 1.**
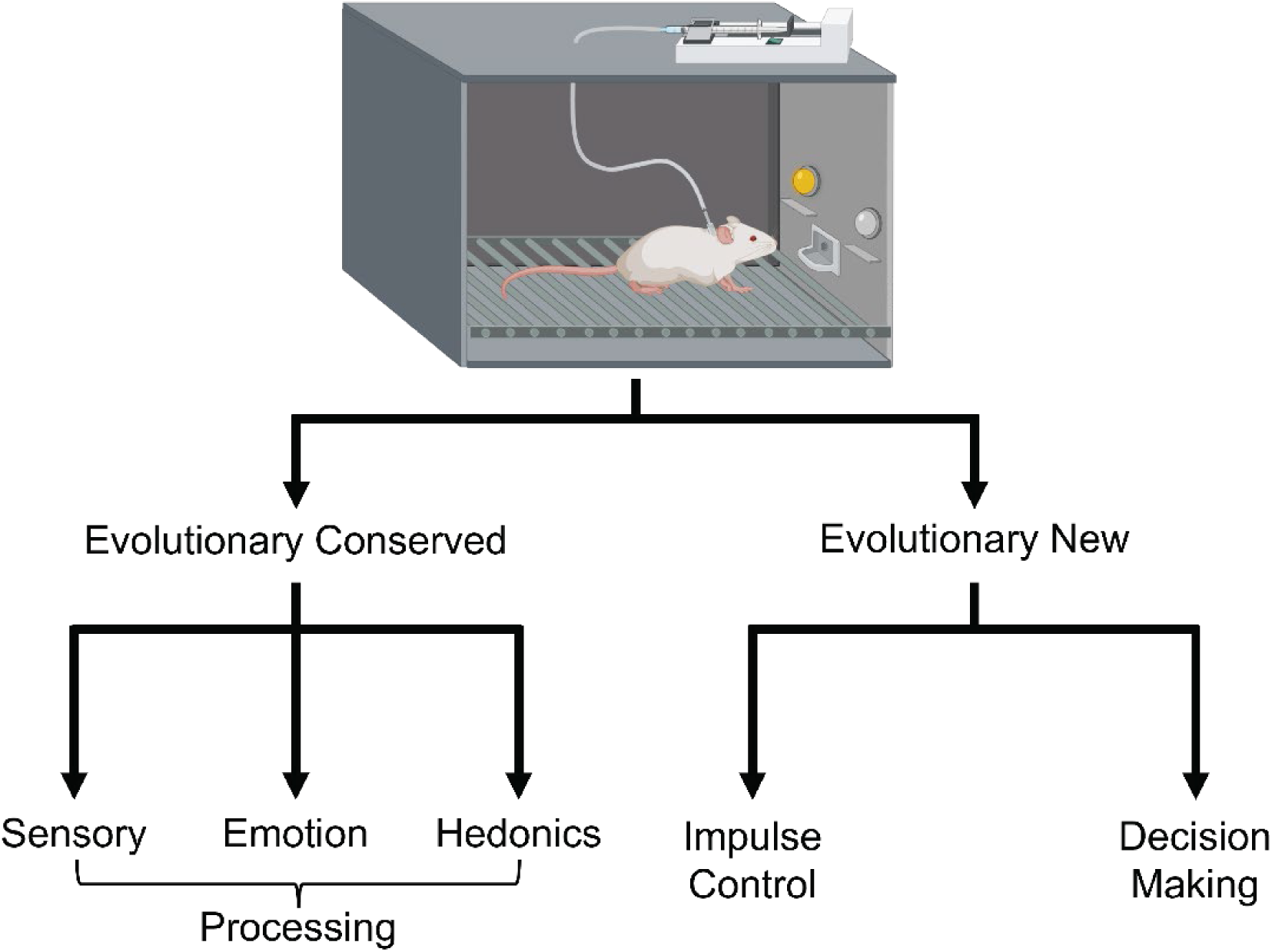
Dissecting drug-seeking behavior into evolutionary-derivedcognitive domains. Drug-seeking behaviorinvolves distinct cognitive processes thatdifferin their evolutionary origins and neurobiological underpinnings. Evolutionarily conserved domains encompass sensory, emotional,and hedonicprocessing,whichare fundamentalfor survival and shared across species. In contrast,evolutionarily newer domains, such as impulse control and decision-making, build upon conserved systems to enable complex cognition unique to phylogeneticallyadvanced species.This frameworkhighlights the distinction between conserved and novel cognitive circuitries,aiding in the identification of specificsignalingcomponents (e.g., receptors and transporters) involved in these behaviors.

### Evolutionary Conserved Behaviors Unaffected by Disrupted Sxc Function

First, visual sensory processing was assessed using the visual discrimination task. Visual discrimination represents a fundamental capability that evolved as vertebrates progressed from basic orientation vision to object vision, enabling detection and classification of objects in complex environments (Nilsson (2022)). This adaptation, conserved across vertebrates but refined in primates, illustrates how early sensory processing systems provide the foundation for discriminative abilities that precede higher cognitive functions (Knudsen (2020)). In this task, rats learned to distinguish between two visual stimuli (airplane and spider), where the positive conditioned stimulus (CS+) yielded a food reward while the negative stimulus (CS-) provided no reinforcement (Fig 2A). MSxc rats did not differ from WT in duration to reach criterion for visual discrimination training (Fig 2B; t_13_ = 0.697, p > 0.05, *d* = 0.361) or discrimination accuracy when achieving criterion (Fig 2C; t_13_ = 0.663, p > 0.05, *d* = 0.343). These findings were not confounded by genotypic differences in associative learning capabilities (Hess et al. (2023)).

**Figure 2.**
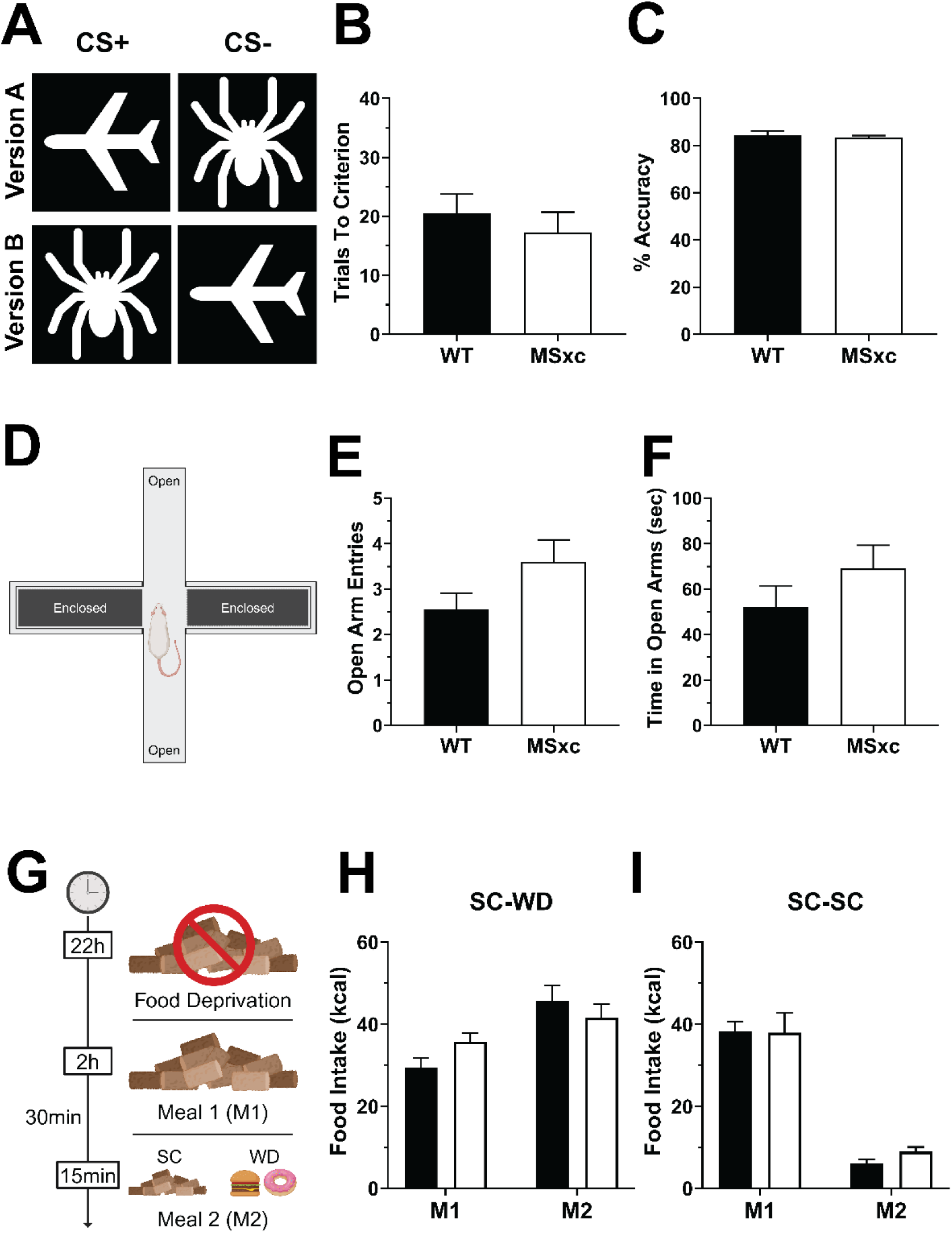
Disrupting Sxc function does not impact evolutionary conserved behaviors. **(A-C)** Visual discrimination tasks using distinct object pairs (airplane and spider) revealed no differences in trials to criterion **(B)** or discrimination accuracy **(C)** between WT and MSxc rats. **(D-F)** Elevated plus maze (EPM) tests showed comparable anxiety-related behaviors, with no significant differences in open arm entries **(E)** or time spent in open arms **(F)**. **(G–I)** In a Two-meal paradigm, MSxc rats displayed normal homeostatic feeding during Meal 1 (M1, standard chow - SC) and hedonic feeding during Meal 2 (M2, SC or western diet - WD), increasing intake when offered WD **(H)** but reducing intake when offered SC **(I)**. 0.001;ηp^2^= 0.233) and MSxc rats (Fig 3B; Meal: F(1,12)= 95.30, p < 0.0001,ηp^2^= 0.553;Meal x Session: F(10,120)= 2.164,p < 0.05,ηp^2^= 0.153) expanded their daily food intake when offered WD, butnot SC,resultingin an increase in body weight (Fig 3C-D; Meal xSession -WT: F(10,120)= 7.580, p < 0.001,ηp^2^= 0.387; MSxc: F(10,120)= 12.530,p <0.0001,ηp^2^= 0.511).

Next, we screened anxiety-based emotional processing using the elevated plus maze (EPM). Anxiety is an evolutionary conserved response that has been maintained for its survival benefits. The elevated plus maze is composed of two closed-arms that shield the rat from light and two open-arms that are exposed to light and surrounding stimuli within the testing environment (Fig 2D). Rats that spend less time in open-arms and execute fewer open-arm entries are considered more anxious (i.e., anxiogenic). To further support our hypothesis that Sxc is dispensable for evolutionary conserved behaviors, MSxc rats displayed no differences in open-arm time (Fig 2E; t_40_ = 1.235, p > 0.05, *d* = 0.382) or entry frequency (Fig 2F; t_40_ = 1.694, p > 0.05, *d* = 0.523) in the EPM, indicating normal anxiety-related, emotional processing.

Lastly, we examined hedonic processing using the two-meal paradigm that separates homeostatic from hedonic drive for feeding behavior. While hedonic-driven behaviors emerged in early bilaterians (vertebrates and invertebrates) through directional adjustment in movement (steering), homeostatic cellular and molecular networks date back to the beginning of life (Torday (2015); Bennett (2021)). In the two- meal paradigm, rats were trained to consume their daily calorie intake of standard chow (SC) in 2-hours (homeostatic feeding; meal 1, M1) until their daily body weight stabilized. Upon stabilization, a brief 15-minute feeding window was introduced shortly after their daily 2-hour feeding bout, offering either the same SC or a highly palatable western diet (WD) (hedonic feeding; meal 2, M2) (Fig 2G). MSxc and WT rats both consumed similar M1 (homeostatic) and M2 (hedonic) meal sizes (Fig 2H; genotype: F_(1,14)_ = 0.175, p > 0.05, η_p_^2^ = 0.009; genotype x meal: F_(1,14)_ = 2.588, p > 0.05, η_p_^2^ = 0.156; M1: WT = 29.38 ± 2.40 kCal, MSxc = 35.75 ± 2.11 kCal; M2: WT = 45.71 ± 3.80 kCal, MSxc = 41.60 ± 3.36 kCal). Furthermore, both genotypes consumed more during M2 when offered WD (Fig 2H; meal: F_(1,14)_ = 11.58, p < 0.05, η_p_^2^ = 0.453; M1: 32.56 ± 1.75 kCal; M2: 38.11 ± 2.50 kCal) illustrating no difference in homeostatic- or hedonic-based feeding behavior. As expected, both MSxc and WT consumed less during M2 when offered SC (Fig 2I; meal: F_(1,10)_ = 163.2, p < 0.05, η_p_^2^ = 0.942; genotype: F_(1,10)_ = 0.188, p > 0.05; η_p_^2^ = 0.030; genotype x meal: F_(1,10)_ = 0.482, p > 0.05, η_p_^2^ = 0.046; M1: WT = 38.33 ± 2.32 kCal, MSxc = 38.00 ± 4.76, Combined = 38.17 ± 2.53 kCal; M2: WT = 6.00 ± 1.10 kCal, MSxc = 9.00 ± 1.10 kCal; Combined = 7.5 ± 0.87 kCal), illustrating intact hedonic assessment of available food resources. Consequently, both WT (Fig 3A; Meal: F_(1,12)_ = 71.60, p < 0.0001, η_p_^2^ = 0.553; Meal x Session: F_(10,120)_ = 3.641, p < 0.001; η_p_^2^ = 0.233) and MSxc rats (Fig 3B; Meal: F_(1,12)_ = 95.30, p < 0.0001, η_p_^2^ = 0.553; Meal x Session: F_(10,120)_ = 2.164, p < 0.05, η_p_^2^ = 0.153) expanded their daily food intake when offered WD, but not SC, resulting in an increase in body weight (Fig 3C-D; Meal x Session - WT: F_(10,120)_ = 7.580, p < 0.001, η_p_^2^ = 0.387; MSxc: F_(10,120)_ = 12.530, p < 0.0001, η_p_^2^ = 0.511).

**Figure 3.**
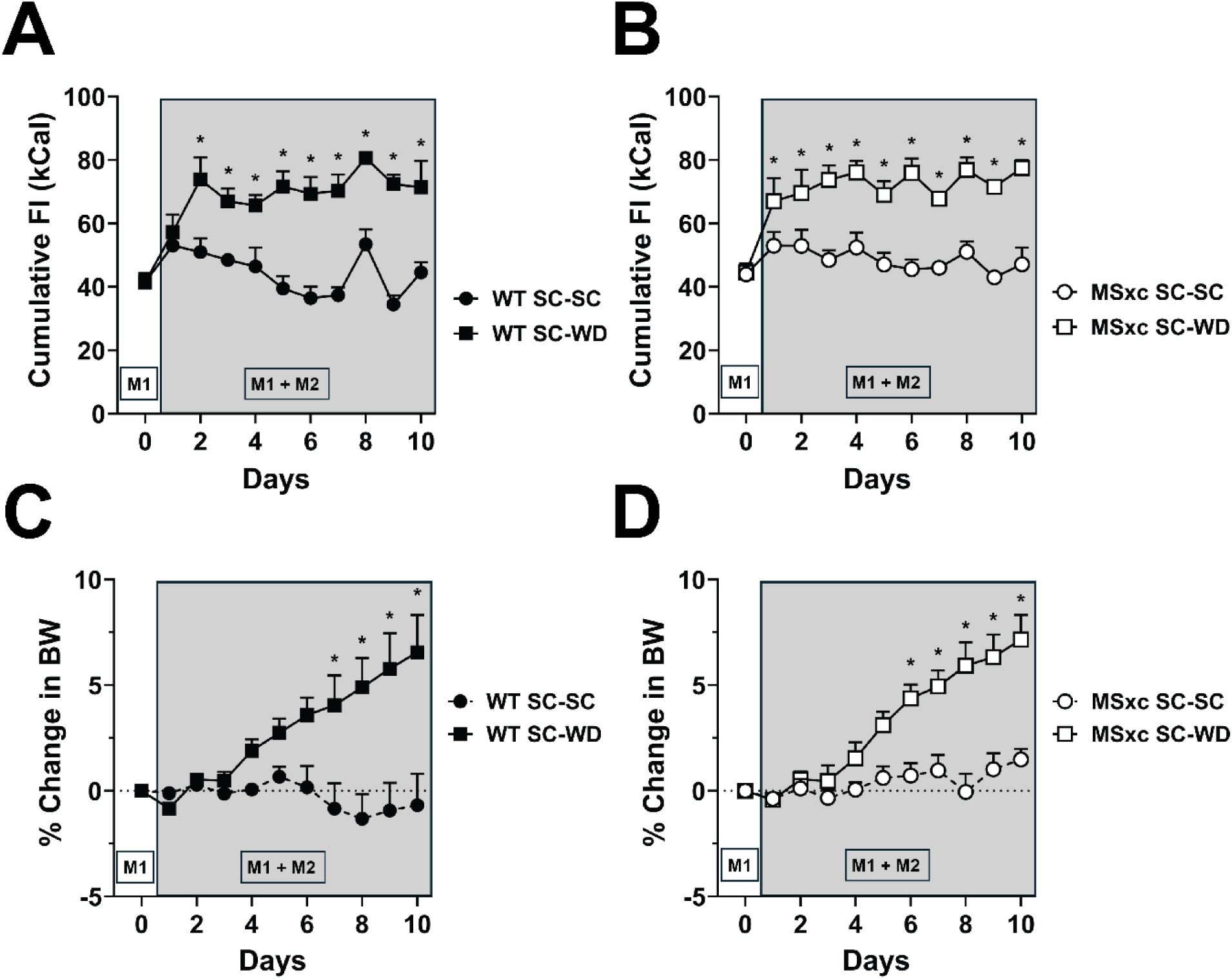
Cumulative food intake (FI) and body weight (BW) changes in theTwo-mealparadigm. After stabilizing cumulative FI during Meal1 (M1) training with standard chow (SC), rats were offered Meal 2 (M2) consisting of eitherSCor a western diet (WD).**(A-B)**Both WT and MSxc rats increased cumulative FI when M2 consisted of WD but maintained stable FI when offered SC.**(C-D)**Consequentially, SC-WD ratsforboth genotypesexhibited significantincreases in body weightduring the M1+M2 phase, while SC-SC rats showed no weight gain.*indicates asignificant effectofmeal type, Holm-Sidakp < 0.05.

**Figure 4.**
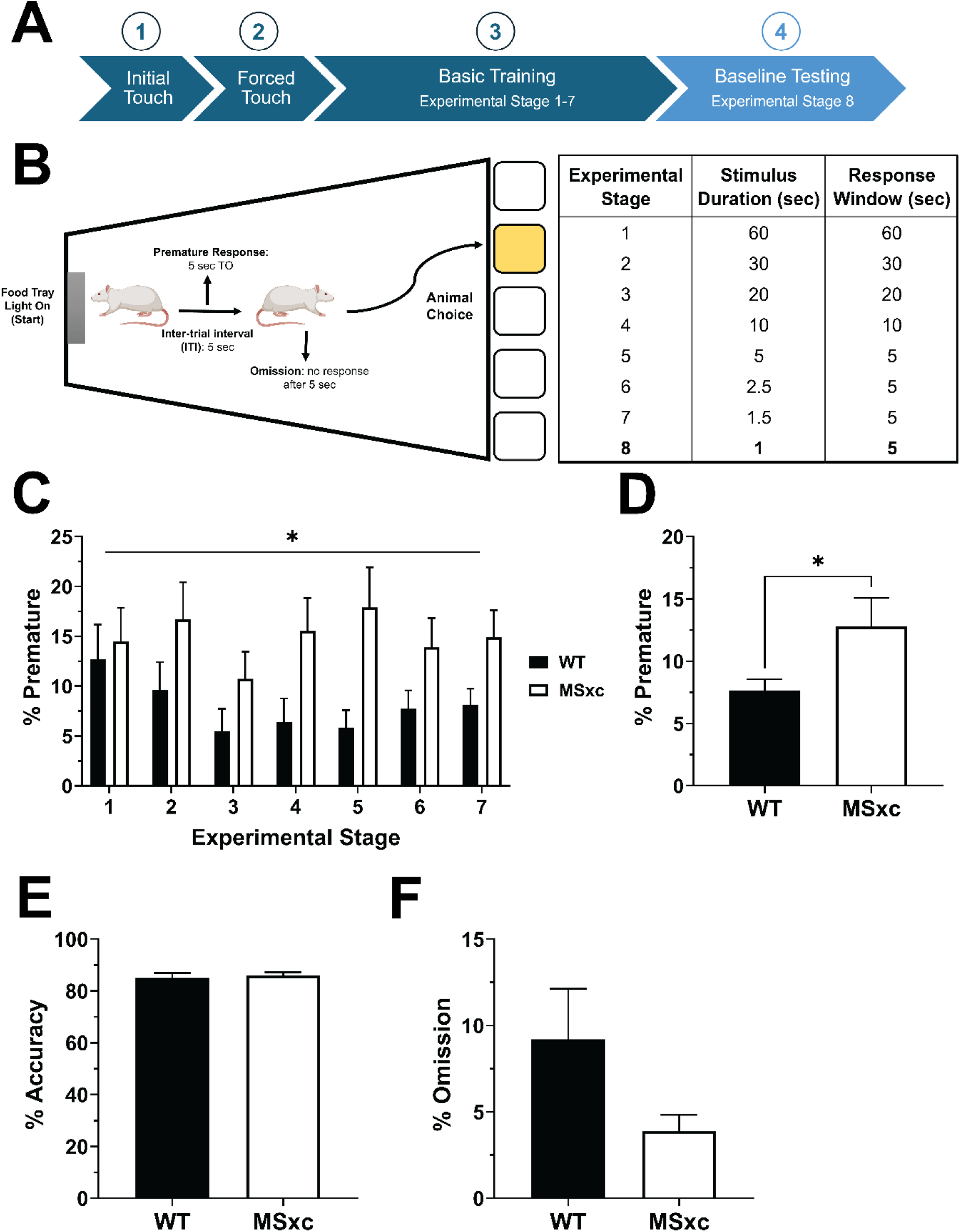
Sxc is required for impulse control, an evolutionary new cognitive function. **(A-B)** Rats were trained in a touchscreen-based 5-choice serial reaction time task to respond (nosepoke) to a light stimulus for a food reward. Training progressed through stages (1–7) with decreasing stimulus durations and response windows, culminating in baseline testing (stage 8). Premature responses (nosepokes before light stimulus appears), response accuracy (nosepoke to light stimulus), and omissions (failure to respond within the response window) were measured. **C)** MSxc rats exhibited higher rates of premature responses compared to WT rats, regardless of training stage difficulty. **D)** On the final day of testing, MSxc rats maintained elevated premature responding. **E-F)** Impulse control deficits in MSxc rats were not attributable to differences in response accuracy **(E)** or task engagement, as omission rates were comparable between groups **(F)**. * indicates significant main effect of genotype, p < 0.05.

**Figure 5.**
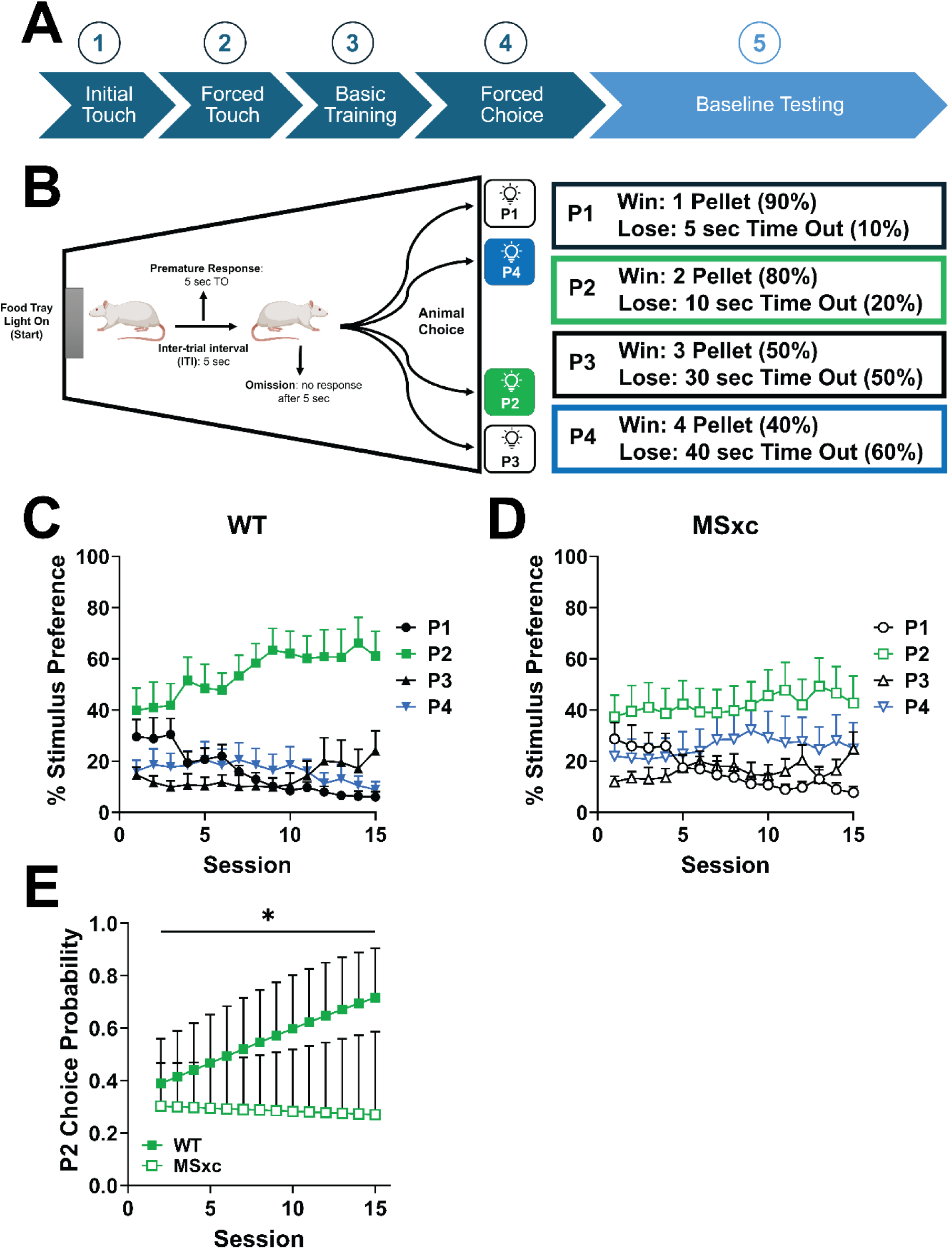
Sxc is required for decision making, an evolutionary new cognitive function. **(A-B)** Rats were trained in a touchscreen-based rat gambling task to develop strategies for maximizing food rewards while minimizing punishment. **A)** Training progressed through stages, culminating in Forced Choice, where rats learned fixed reward and punishment schedules for each stimulus (P1–P4). **B)** Stimuli differed in reward density, win probability, and punishment duration (Time Out). **(C-D)** WT rats identified P2 as the optimal stimulus, increasing preference over sessions, while MSxc rats failed to develop this strategy. **E)** By the final session, WT rats primarily selected P2 (72%), whereas MSxc rats chose P2 at chance levels (27%). * indicates significant genotype × session interaction, GLMM p < 0.05.

**Figure 6.**
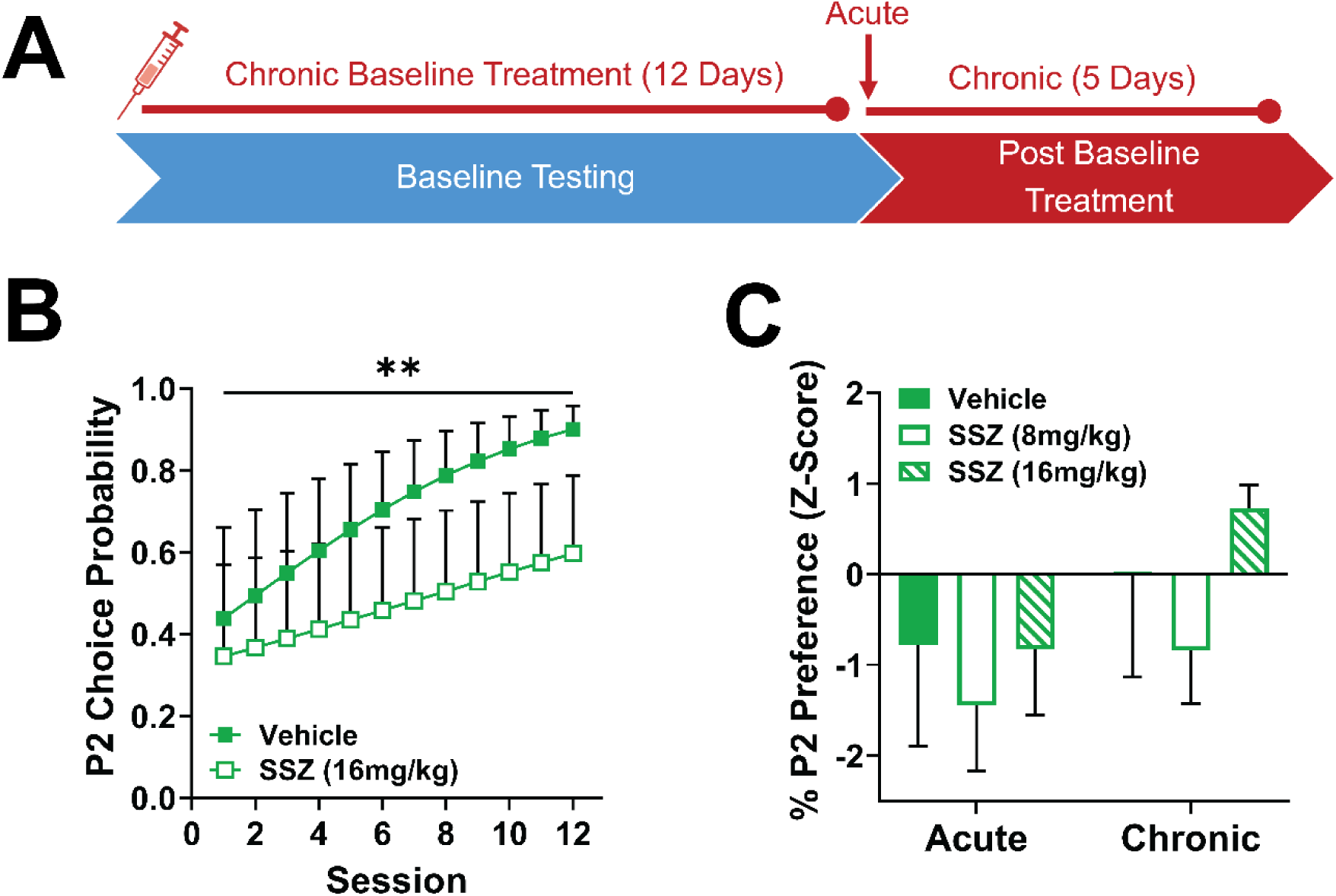
Sxcis required for optimal strategy development but not for maintaining learnedstrategies. **A)**WT rats were trained in the rat gambling task and treated with vehicle or the Sxcinhibitor sulfasalazine (SSZ) during either strategy development (ChronicBaseline Treatment) or afterstrategies were established (Acute:1 session; Chronic: 5 sessions).**B)**During baseline testing,WTrats treated with a high dose ofSSZ (16 mg/kg) showed impaired selection of the optimal stimulus(P2), as indicated by reduced choice probability compared to vehicle-treated rats.**C)**In contrast,acute and chronicSSZ treatment (8 mg/kg or16 mg/kg) following strategy establishment did notaffectP2 preference, demonstrating thatSxc is dispensable for maintaining learned strategies.** indicatessignificant treatment x session interaction, GLMM p <0.01.

Collectively, these results indicate that evolutionary conserved behaviors were maintained following functional elimination of Sxc (MSxc), recapitulating previous findings of unaffected basic brain function (e.g., general activity patterns in open field and basic memory in novel object recognition task) (Hess et al. (2023)). Therefore, we sought to assess the functional necessity of Sxc in evolutionary new behaviors.

### Disrupting Sxc Function Produces Deficits in Evolutionary New Behaviors

Next, we assessed the necessity of Sxc in impulse control, a cognitive function crucial for higher-order processing in evolutionarily advanced species (Kaczanowska et al. (2022)). While impulse control exists across diverse species, its neural implementation varies significantly. For instance, some birds demonstrate sophisticated impulse control despite lacking a prefrontal cortex (PFC), instead developing this ability independently through distinct neural structures and pathways (Emery and Clayton (2004); Kabadayi et al. (2016)). In mammals, however, impulse control capabilities correlate with PFC expansion, enabling more flexible inhibitory control (Maclean et al. (2014); Isaksson et al. (2018)). Notably, PFC expansion in primates and humans coincided with increased astrocyte complexity and diversity, potentially enhancing Sxc’s involvement in information processing (Oberheim et al. (2009); Vasile et al. (2017)).

Here, we inspected impulse control using the 5-choice serial reaction time task (5CSRT). Rats were trained to associate a light stimulus with a reward (Initial Touch), engage with the light stimulus to receive a reward (Forced Touch), and increase their response latency towards the light stimulus (Basic Training) (Fig 4A-B). As stimulus duration shortened, rats were forced to respond more quickly and develop a prepotent response – enhancing the urge to engage with the light stimulus. We found that regardless of training difficulty, MSxc rats exhibited heightened premature responding compared to WT rats (Fig 4C; genotype: F_(1,11)_ = 5.608, p < 0.05, η_p_^2^ =0.378; genotype x experimental stage: F_(6,66)_ = 1.226, p > 0.05, η_p_^2^ = 0.101). Premature responding did not increase with training difficulty (Fig 4C; experimental stage: F_(6,66)_ = 1.566, p > 0.05, η_p_^2^ = 0.125), indicating that training stages adequately challenged impulse control without provoking anticipatory responses. Consistent with training results, baseline testing revealed elevated premature responding in MSxc rats compared to WT (Fig 4D; t_11_ = 2.217, p < 0.05, *d* = 1.233). Interestingly, MSxc displayed similar response accuracy and omission during baseline testing (Fig 4E-F; accuracy: t_11_ = 0.356, p > 0.05, *d* = 0.198; omission: t_11_ = 1.605, p > 0.05, *d* = 0.893), indicating functional working memory and no change in task engagement, respectively.

Next, we measured decision making using the rat gambling task (rGT). Decision making abilities evolved in response to ecological challenges. Advanced species navigate these complexities through enhanced inhibitory control mechanisms in evolutionary recent structures like the prefrontal cortex. These mechanisms optimize cognition by filtering information to influence choices and enable adaptation to changing conditions. In complex tasks like the rGT, which integrates multiple cognitive demands, astrocytes may selectively activate specific neuronal patterns appropriate to changing task conditions (Murphy-Royal et al. (2023)). These task-specific adaptations orchestrated by astrocytes may be facilitated through Sxc.

In the rGT, like the 5CSRT task, rats were sequentially trained to engage with a light stimulus positioned on the touchscreen and learned to associate stimulus position with a designated reward/punishment schedule that varied in probability, reward density, and timeout duration (Fig 5A-B; Forced Choice). During baseline testing, rats simultaneously encountered all stimuli – free to select any option. Stimulus selection percentages were recorded in 30-minute sessions across 15 days to determine stimulus preference as a metric for optimal decision making. Based on varying stimulus schedule reward densities, win probabilities, and timeout durations, each stimulus (if selected in isolation) would result in different reward totals by the end of the session: P1, 295; P2, 411; P3, 135; and P4, 99 (Kim et al. (2017)). Therefore, allocation of stimulus selection towards P2 would indicate optimal strategy execution in maximizing reward outcomes.

MSxc rats exhibited impaired decision-making relative to WT rats, showing little or no development of preference for the optimal (P2) stimulus across sessions (Fig 5C– D). This difference was confirmed by generalized linear mixed model (GLMM) analysis. A significant main effect of initial preference (earlypref) was found (Type III Wald χ²_(2)_ = 29.63, *p* < 0.0001), confirming effectiveness of the baseline preference categorization. However, initial preference did not significantly interact with genotype or session (all interactions, p > 0.64) indicating its effects remained consistent across groups and sessions. Importantly, a significant Session x Group interaction (*X*^2^_(1)_ = 3.96, p < 0.05) demonstrated differential learning trajectories between genotypes: WT rats significantly increased their preference for the optimal stimulus (P2) across sessions, whereas the MSxc rats failed to increase optimal (P2) choice behavior across testing. By the final test session (Session 15), WT rats selected the optimal stimulus (P2) 72% of trials, whereas MSxc rats remained at chance levels (27%) (Fig 5E; genotype: *X*^2^_(1)_ = 0.123, p > 0.05; session: *X*^2^_(1)_ = 0.008, p > 0.05; genotype x session: *X*^2^_(1)_ = 4.171, p < 0.05).

To further validate the critical role of Sxc in optimal decision-making, we employed a complementary pharmacological approach. WT rats were trained up to the testing phase and separated into treatment groups using a randomized block design based on pre-treatment performance during the forced choice training phase. Rats received either vehicle or sulfasalazine (SSZ), a selective inhibitor of Sxc previously shown to produce deficits in cognitive flexibility (Lutgen et al. (2014)). Rats were systemically treated with SSZ at a high dose (16mg/kg, IP, 2 hrs before daily testing) for 12 days of testing (Fig 6A, Chronic Baseline Treatment).

Rats treated chronically with SSZ exhibited impaired development of optimal stimulus selection (P2) across testing sessions relative to vehicle-treated controls, as confirmed by generalized linear mixed model (GLMM) analysis (Fig 6). The Type III Wald chi-square tests indicated a significant main effect of Session (χ²_(1)_ = 6.97, *p* = 0.008), reflecting that, overall, rats showed improvements in P2 selection over time. Importantly, a significant Session × Treatment interaction (χ²_(1)_ = 6.64, *p* = 0.010) indicated differential learning trajectories: Vehicle-treated rats progressively increased their estimated optimal P2 choice probabilities across sessions significantly above chance level starting as early as session 2 (49%, 95% CI: 0.29-0.70), whereas SSZ-treated rats failed to perform above chance level until session 6 (45%, 95% CI: 0.27-0.66). Notably, both Vehicle- and SSZ-treated rats continued to significantly exceed chance (25%) at session 12, but SSZ-treated rats remained significantly impaired relative to Vehicle controls, with estimated optimal-choice probabilities of 90% (95% CI: 0.78–0.96) and 60% (95% CI: 0.37-0.79) respectively. Further complexity in these effects was evidenced by a significant three-way Session × Treatment × Version interaction (χ²_(1)_ = 3.87, *p* = 0.049), suggesting that the position of the optimal stimulus (Version A vs. B) interacted with treatment effects over time. No significant main effects were observed for Treatment (χ²_(1)_ = 0.15, *p* = 0.70) or Version alone (χ²_(1)_ = 0.49, *p* = 0.48). Taken together, on the final test (session 12), vehicle-treated rats demonstrated enhanced preference for the optimal stimulus (P2), whereas SSZ-treated rats’ performance remained impaired (Fig 6B).

To determine if there was a temporal dependency of Sxc function in optimal decision making, we investigated optimal strategy maintenance after the baseline learning period. We trained WT rats through baseline testing and then split them into vehicle and two SSZ groups (low dose: 8mg/kg; high dose: 16mg/kg) to assess two timepoints of Sxc inhibition: acute (1 session) and chronic (5 sessions) (Fig 6A; Post-Baseline Treatment). We accounted for individual baseline variability of P2 preference prior to SSZ treatment using z-score standardization on the final three days of baseline testing (session 10-12). Inhibition of Sxc did not affect P2 preference following strategy development (Fig 6C; treatment: F_(2, 21)_ = 0.311, p > 0.05, η_p_^2^ = 0.039; timepoint: F_(2,42)_ = 2.109, p > 0.05, η_p_^2^ = 0.091; treatment x timepoint: F_(4, 42)_ = 0.322, p > 0.05, η_p_^2^ = 0.030). These results indicate that Sxc plays an essential role in strategy development for optimal decision making but is not required to maintain these strategies.

## Discussion

This study demonstrates a restricted role of system xc- (Sxc) in advanced cognitive functions associated with evolutionary recent species. We isolated components of basic (sensory, emotional, and hedonic processing) and higher-order cognition (impulse control and decision-making) to elucidate their discrete or synergistic contributions to pathological behaviors, such as cocaine use disorders (CUDs).

Previous findings suggest that Sxc, the cystine-glutamate antiporter expressed primarily on astrocytes (Ottestad-Hansen et al. (2018)), is an integral molecular component in controlling drug seeking behavior (Baker et al. (2002); Baker et al. (2003), Hess et al. (2023)). We implemented a comprehensive cognitive battery to investigate the role of Sxc in both evolutionarily conserved and more recently evolved complex behaviors that may drive drug seeking.

First, we measured simple, evolutionary conserved cognitive processing systems: sensory, emotional, and hedonics. Genetically modified rats that lack Sxc function (MSxc) show intact basic, evolutionary conserved processing capabilities in the visual discrimination task (sensory), elevated plus maze (emotional), and Two-Meal paradigm (hedonic). Sxc is an evolutionary recent mechanism that emerged following the divergence of deuterostomes from protostomes approximately 550 million years ago (Erwin and Davidson (2002); Lewerenz et al. (2013)). Despite this divergence, genetic patterning mechanisms organizing brain development remained highly conserved between these lineages (Hirth et al. (2003); Arendt et al. (2016)). This suggests that Sxc’s emergence did not coincide with fundamental changes in brain architecture that are often aligned with cognitive enhancement (Deaner et al. (2007); Beaulieu-Laroche et al. (2021)). Rather, Sxc likely provided novel signaling capabilities that were subsequently recruited during vertebrate evolution and refined within mammalian phylogenies, millions of years after the initial protostome-deuterostome split, to support more complex cognitive functions. The evolutionary conserved cognitive processes measured in this study predate this evolutionary innovation, explaining why they remain intact despite Sxc loss.

In the visual discrimination task, we found that MSxc rats did not differ from WT in the discrimination between two distinct object pairs (airplane and spider) to obtain a food reward. This preserved ability in MSxc rats suggests that basic visual discrimination relies on neural mechanisms that evolved prior to Sxc emergence. Visual processing exemplifies how fundamental sensory systems, rooted in evolutionary preserved circuitries, have been refined within more advanced species to accommodate their more sophisticated ecological niches. Visual processing initially evolved as non-directional photoreception for basic functions like circadian entrainment, later developing into directional photoreception, and eventually refining into ancient vision systems capable of shape discrimination – a fundamental ability we specifically assessed in this study (Nilsson (2009); Nilsson et al. (2014); Nilsson (2022)). In more advanced species, these ancient pathways have been elaborated upon, ultimately integrating object vision with higher-order cognitive processes for enhanced environmental and situational awareness, such as visual perception. Ultimately, incremental evolutionary refinement of basic sensory mechanisms, including vision, emotion, and hedonics, provided a stable neural foundation upon which more complex cognitive abilities could be built, allowing organisms to adapt to increasingly complex ecological niches (Anderson and Adolphs (2014); Berridge and Kringelbach (2015); Knudsen (2020)).

Our findings in the elevated plus maze (EPM) paradigm indicate that MSxc rats exhibit similar anxiety-related behaviors to wild-type (WT) rats. This result is significant because it suggests that the genetic disruption of Sxc does not inherently augment anxiety, which could otherwise confound behavioral deficits observed in other paradigms. This is noteworthy given that disruptions in oxidative stress maintenance, such as those affecting glutathione production, have been linked to anxiogenic behaviors (Hovatta et al. (2005); Masood et al. (2008); Patki et al. (2013)). Notably, redox maintenance is an evolutionary ancient process essential for life, yet eliminating Sxc function, which disrupts glutathione production, did not alter anxiety-related behavior in our model. The lack of anxiogenic effects in MSxc rats highlights that alternative, degenerate antioxidant mechanisms may compensate for the absence of Sxc, potentially altering stress sensitivity or homeostatic setpoints without impacting anxiety.

The Two-Meal paradigm revealed that hedonic processing is preserved despite disrupting Sxc, which encompasses both *liking* (pleasure derived from rewards) and *wanting* (motivation to pursue rewards) (Zheng and Berthoud (2007); Berridge et al. (2009)). Both WT and MSxc rats exposed to western diet (WD) during Meal 2 displayed similar elevations in food intake compared to the stable consumption levels seen in standard chow (SC) controls, reflecting intact *liking*. This behavior aligns with the role of opioid and endocannabinoid signaling within hedonic hotspots, such as the nucleus accumbens (NAcc), ventral pallidum (VP), and orbitofrontal cortex (OFC), which mediate the pleasurable aspects of reward (Jarrett et al. (2005); Smith et al. (2011); Morales and Berridge (2020)). Additionally, the progressive increase in body weight across sessions for WD-fed rats highlights sustained motivation to eat (*wanting*), even in the face of potential satiety. The weight gain in WD-fed rats may also result from altered homeostatic food intake setpoints through allostasis, a hypothesis that requires further experimental validation. Together, these findings verify that Sxc disruption does not impair the degenerate neural systems underlying hedonic processing. This ensures that behavioral deficits observed in complex cognitive tasks are not confounded by impairments in fundamental mechanisms of *liking* or *wanting*.

Next, we characterized MSxc rats across more advanced, evolutionary new behavioral features that are essential for higher-order cognitive processing. First, we measured impulse control using the 5-choice serial reaction time task (5CSRT). Deficits in impulse control, known as impulsivity, are a hallmark trait in neuropsychiatric disorders, ranging from substance abuse to Parkinson’s disease (de Wit (2009); Kozak et al. (2019); Vassileva and Conrod (2019); Nombela et al. (2014)). The nuance of impulsivity makes it a challenging phenotype to characterize, manifesting as impaired motor control (impulsive action) or decision making without foresight (impulsive choice); both of which differ in their underlying cognitive circuitries (Dalley et al. (2011), Nautiyal et al. (2017), Dalley and Robbins (2017)). The 5CSRT task specifically measures impulsive action through inhibitory control over prepotent responses, where deficits in this control result in premature actions. MSxc rats exhibit elevated premature responding across training and testing, but did not differ from WT in working memory (response accuracy) or task engagement (omission). It is possible that the observed impulsive actions in MSxc rats were caused by elevated incentive salience of the light stimulus, making resisting a prepotent response more challenging. Previous evidence found that MSxc rats exhibit increased sign-tracking in pavlovian conditioning and higher levels of drug seeking behavior in a cocaine reinstatement paradigm (Hess et al. (2023)). Behavioral outcomes representative of incentive salience could also be caused by the weakening of response inhibition, which is an example of phylogenetically enhanced cognitive control seen in humans and other advanced species (for response inhibition: Flagel et al. (2008); Tomie and Morrow (2018); Sarter and Phillips (2018); Anselme and Robinson (2020); for phylogenetic cognitive control: Ardila (2008); Grant (2016); Dukas (2017); Van Duijn (2017)). In support of this, MSxc rats did not differ from WT in acquisition, maintenance, and extinction of cocaine-reinforced operant respond, arguing against the incentive salience hypothesis (Hess et al. (2023)). These findings illustrate the discrete nature of Sxc’s role in evolutionary new forms of cognition, such as impulse control, without producing more diffuse deficits that impact working memory or overall task engagement.

Lastly, we investigated Sxc’s role in decision making within the rat gambling task (rGT). In this preclinical analog of the Iowa Gambling Task (IGT), rats must integrate complex information spanning reward probabilities, loss frequencies, and punishment duration to develop an optimal decision-making strategy that maximizes rewards earned. The multifaceted nature of this integration makes this task particularly challenging and sensitive to subtle cognitive deficits. We found that MSxc rats were incapable of developing and executing optimal decision-making strategies, exhibited by the lack of optimal stimulus (P2) selection above chance (25%) across sessions.

To ensure MSxc deficits specific to Sxc dysfunction rather than developmental compensatory mechanisms, we used the Sxc inhibitor sulfasalazine (SSZ). SSZ has been shown to impair cognitive flexibility (Lutgen et al. (2014)) without causing widespread cognitive deficits (Verbruggen et al. (2021)), highlighting its targeted effects. SSZ treatment recapitulated the MSxc phenotype when administered during strategy development (Chronic Baseline Treatment) but not after decision-making strategies were established (Post-Baseline Treatment: Acute and Chronic). While SSZ-treated rats increased their preference for P2 across sessions, their rate of improvement was significantly slower than vehicle controls, differing from MSxc rats, whose P2 preference remained at chance levels. This discrepancy may stem from: 1) SSZ’s metabolism allowing partial Sxc recovery within 24 hours, enabling consolidation of learned events; or 2) prior exposure to reward/punishment schedules during training without Sxc inhibition, allowing some strategy development before testing. Future studies could address this by administering SSZ during training to better mimic the MSxc experience.

## Conclusion

Glutamate is one of the most ubiquitous neurotransmitters in the brain, orchestrating processes that range from basic sensory perception to highly complex cognitive functions. Its central role in nearly all forms of brain activity makes it an enticing therapeutic target for neuropsychiatric disorders. However, the molecular degeneracy of glutamate signaling – where overlapping pathways mediate a wide range of functions – poses significant challenges for developing targeted therapies without inducing widespread side effects. In this context, system xc- (Sxc) emerges as a promising solution. As an evolutionary new signaling protein within the glutamate system localized to astrocytes, Sxc deviates from canonical glutamate signaling by selectively influencing advanced cognitive domains while sparing fundamental conserved processes. Sxc represents an exciting frontier in personalized medicine, offering the potential to revolutionize treatments for neuropsychiatric disorders by addressing discrete cognitive deficits with unprecedented precision and safety.

## Data availability statement

The data that support the findings of this study are available from the corresponding author upon reasonable request.

## Funding statement

This work was supported by the National Institute on Drug Abuse of the National Institutes of Health under Award Number R01DA050180, the National Center for Advancing Translational Sciences under Award Number TL1TR001437, and grants from the JJ Keller Foundation and the Charles E. Kubly Mental Research Center.

## Conflict of interest disclosure

DAB is the co-founder of, and owns shares in, Promentis Pharmaceuticals. Promentis is developing glutamatergic compounds to treat impulse control disorders but was not involved in these studies.

## Ethics approval statement

The experimental protocol involving animals were ethically reviewed and approved in accordance with guidelines for the care and use of laboratory animals, MU IACUC (AR-3815).

## Notes

### Competing Interest Statement

The authors have declared no competing interest.

